# OPUS-GO: An interpretable protein/RNA sequence annotation framework based on biological language model

**DOI:** 10.1101/2024.12.17.629067

**Authors:** Gang Xu, Ying Lv, Ruoxi Zhang, Xinyuan Xia, Qinghua Wang, Jianpeng Ma

**Affiliations:** Multiscale Research Institute of Complex Systems Fudan University Shanghai, 200433, China; Shanghai AI Laboratory Shanghai, 200030, China; Fudan University Shanghai, 200433, China; Center for Biomolecular Innovation, Harcam Biomedicines, Shanghai, 200131, China

## Abstract

Accurate annotation of protein and RNA sequences is essential for understanding their structural and functional attributes. However, due to the relative ease of obtaining whole sequence-level annotations compared to residue-level annotations, existing biological language model (BLM)-based methods often prioritize enhancing sequence-level classification accuracy while neglecting residue-level interpretability. To address this, we introduce OPUS-GO, which exclusively utilizes sequence-level annotations to provide both sequence-level and residue-level classification results. In other words, OPUS-GO not only provides the sequence-level annotations but also offers the rationale behind these predictions by pinpointing their corresponding most critical residues within the sequence. Our results show that, by leveraging features derived from BLMs and our modified Multiple Instance Learning (MIL) strategy, OPUS-GO exhibits superior sequence-level classification accuracy compared to baseline methods in most downstream tasks. Furthermore, OPUS-GO demonstrates robust interpretability by accurately identifying the residues associated with the corresponding labels. Additionally, the OPUS-GO framework can be seamlessly integrated into any language model, enhancing both accuracy and interpretability for their downstream tasks.

## Introduction

In recent years, there have been significant breakthroughs in the development of large language models in natural language processing (1–4). By treating the “building blocks” of biological sequences, such as residues in proteins (5) and nucleotides in RNAs (6), as “tokens”, researchers have successfully applied language modeling techniques to the field of biological sequences (5–16). Furthermore, the rapid expansion of biological data has significantly facilitated the development of large-scale biological language models, enabling them to accurately detecting intricate patterns and functional relationships within these biological molecules.

Protein language models (PLMs) have emerged as powerful tools for predicting protein structures, variant effects, and various functional attributes (5, 11). Proteins, consisting of a linear sequence of amino acids, exhibit a specific arrangement that dictates their three-dimensional structure and, ultimately, their functional properties. By training on extensive datasets of protein sequences, PLMs acquire foundational evolutionary knowledge about protein structure and function, thereby enabling them to perform a wide range of protein modeling and functional prediction tasks (17–20). Among PLMs, ESM-2 (5) stands as the most prominent and widely utilized. Based on its features, the end-to-end structure prediction method, ESMFold, has exhibited remarkable performance with a significantly shorter inference time compared to AlphaFold2 (21). Furthermore, its applicability has been demonstrated in numerous other contexts, including protein function prediction, EC number prediction, and various other tasks (18, 20, 22).

Significant advancements have also been made in the development of RNA language models (RLMs), despite the unique challenges posed by the limited four-letter alphabet and the relatively lower conservation of RNA sequences compared to proteins (15, 23). These challenges can result in sparsity and noise within RNA data, thereby complicating the development of RLMs (6). To address these issues, researchers have recently introduced various pretraining strategies, such as a multilevel masking strategy (16) and a protein-derived transfer learning strategy (6). These strategies have enabled RLMs to more effectively extract evolutionary and co-evolutionary information, leading to substantial improvements in tasks associated with RNA structural and functional predictions (6, 14, 16). Among the recently released RLMs, RiNALMo (14) and ProtRNA (6) are two notable methods. RiNALMo has been trained on a dataset consisting of 36 million non-coding RNAs (ncRNAs), incorporating architectural modifications inspired by Llama2 (24), such as rotary positional embedding and the SwiGLU activation function. In contrast, ProtRNA has been developed using a cross-modality transfer learning strategy, utilizing the parameters borrowed from the ESM-2 model as its initialization. Both models have demonstrated commendable performance in various RNA downstream tasks, including protein-RNA interaction prediction and RNA secondary structure prediction.

In computational biology, interpretability is a critical consideration due to its ability to provide significant biological insights that transcend mere accuracy improvements. This necessity arises from the requirement for biologists to compare and contrast model predictions, and to integrate these predictions with experimental data and theoretical insights. However, the guidelines for employing interpretable machine learning in computational biology remain largely underdeveloped (25). Given the successful utilization of features derived from biological language models in addressing biological tasks, the development of an interpretable framework for annotating protein/RNA sequences, based on these features, becomes a pressing concern.

Recently, some researches have introduced the Gradient-weighted Class Activation Map (Grad-CAM) (26) to identify residues within a protein structure that are crucial for predicting specific functions. Grad-CAM is a class-discriminative localization technique that offers visual explanations for predictions made by convolutional neural network (CNN)-based models (27). However, Grad-CAM is a post-hoc method and can only be employed as a post-training analysis tool, providing model interpretability without improving model accuracy. Consequently, we have chosen to develop OPUS-GO within the multiple-instance learning (MIL) framework, which not only offers robust interpretability but also has the potential to concurrently enhance model performance. Moreover, by utilizing the MIL framework, we can obtain an instance-level classifier that can be employed independently of its surrounding context, unlike GAM-based methods that necessitate context-specific application. Therefore, the development of a MIL-based method may avoid disconnection from the training process, thereby exhibit strong generalizability and facilitate broader application.

In this study, we introduce OPUS-GO, an interpretable framework designed to annotate protein/RNA sequences using features derived from the biological language models. Traditionally, in sequence-level prediction tasks, researchers have often utilized the average value of the entire sequence’s features as the representative for the corresponding protein/RNA (6, 8, 14, 28). In contrast, OPUS-GO adopts a setting inspired by the MIL strategy. Specifically, we conceptualize the features of a particular entire sequence as a bag (i.e. sequence-level), with the feature of each residue within it serving as an instance (i.e. residue-level). Note that, we possess only the sequence-level label for each sequence data. By utilizing our modified MIL framework, we are able to train a residue-level classifier that can classify each residue according to the sequence-level label and identify the most representative residues for each corresponding label.

We evaluate the performance of OPUS-GO on 8 protein multi-class or multi-label classification downstream tasks utilizing features derived from ESM-2, and on two RNA downstream tasks employing features derived from RiNALMo and ProtRNA, respectively. The results show that OPUS-GO demonstrates superior classification accuracy compared to its baseline methods for the majority of protein and RNA downstream tasks. Notably, OPUS-GO satisfactorily identifies the residues associated with the corresponding labels. In most tasks, when utilizing the average values of features from three randomly selected residues, a significant decline in performance is observed compared to using the average values of features from the entire sequence. However, when employing the average values of features from three residues selected by OPUS-GO, the accuracies remain relatively stable, highlighting OPUS-GO’s effectiveness in selecting residues associated with the corresponding label. Furthermore, OPUS-GO exhibits effectiveness on datasets with over 5,000 labels, suggesting its broad applicability across diverse scenarios.

## Results

### Assessment of OPUS-GO’s Effectiveness on the MNIST Dataset

To assess the effectiveness of OPUS-GO, we conduct experiments utilizing the MNIST dataset (29), with a multi-label classification task setting (Figure 1a). In this context, individual samples from the MNIST dataset are treated as instances, and are grouped to form bags, with each bag containing 200 instances. Initially, labels to be included in each bag are randomly selected. Then, images belonging to the selected labels are randomly chosen to form the bags. Each bag is assigned a label represented by a 10-class vector of multiple labels, indicating the presence or absence of corresponding images within the bag.

**Figure 1.**
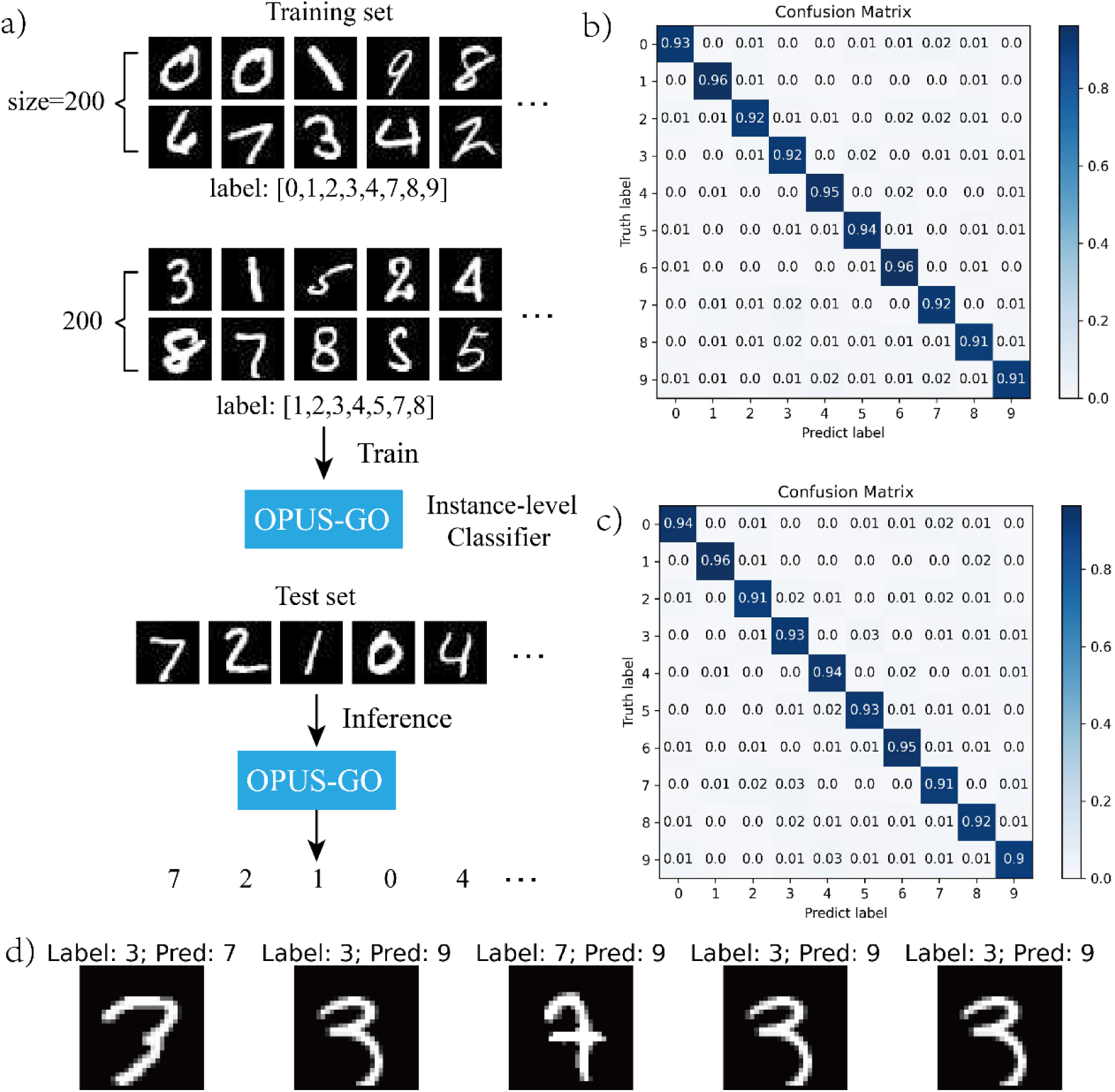
The results of OPUS-GO on the MNIST Dataset. a) The workflow for the multi-label classification task setting. In this setting, each bag contains 20 images, and the label for each bag is represented as a 10-class vector of multiple labels, indicating the presence or absence of the corresponding types of images within the bag. The input features are obtained by reshaping each image to a 28x28-pixel vector. b) The confusion matrix for the results of OPUS-GO on the training set. c) The confusion matrix for the results of OPUS-GO on the test set. d) Several cases where the predictions made by OPUS-GO are incorrect.

As shown in Figure 1b-d, OPUS-GO exhibits the ability to achieve instance-level classification using only bag-level labels with a relatively high degree of accuracy. This suggests that OPUS-GO is effective in identifying instances that associated with their respective bag-level labels, thereby demonstrating its interpretability in the context of the multi-label classification task.

### Performance of OPUS-GO on three protein GO term prediction datasets

The protein function annotation task is aimed at assigning multiple functional labels to a protein. To evaluate the performance of our method, we utilize three standard benchmarks for Gene Ontology (GO) term prediction (27, 30), specifically, molecular function (denoted as GO-MF) with 489 labels, biological process (denoted as GO-BP) with 1943 labels, and cellular component (abbreviated as GO-CC) with 320 labels. All of them belong to the multi-label classification tasks, and we adopt Area Under the Precision-Recall Curve (AUPR) and Fmax as evaluation metrics for each method.

For protein downstream tasks, we utilize features derived from ESM-2, a prominent and widely-adopted protein language model. As shown in Table 1, OPUS-GO demonstrates superior sequence-level classification accuracy in most cases compared to the baseline method ESM-2 (ESM-2 in Table 1), which utilizes the average value of ESM-2 features across all residues as the sequence-level representative. Furthermore, we present the results of several other leading methods. HEAL (31) is grounded in a hierarchical graph Transformer, integrating features derived from the protein language model ESM-1b and coordinates from the corresponding PDB structure for each residue to achieve the final prediction. GGN-GO (32) leverages a geometric graph network to enrich feature extraction by capturing multi-scale geometric structural features at both atomic and residue levels, utilizing features from language models ESM-2 and ProtTrans, as well as structural features. ProtST (33) and PROTLLM (34) are two models that incorporate information derived from natural language models, utilizing additional protein-text data to fine-tune the features from the biological language model ESM-2 for enhanced representation. The results show that OPUS-GO also outperforms these methods in most cases. It is noteworthy that, in contrast to the latter baselines, OPUS-GO and ESM-2 solely use features derived from ESM-2.

**Table 1.**
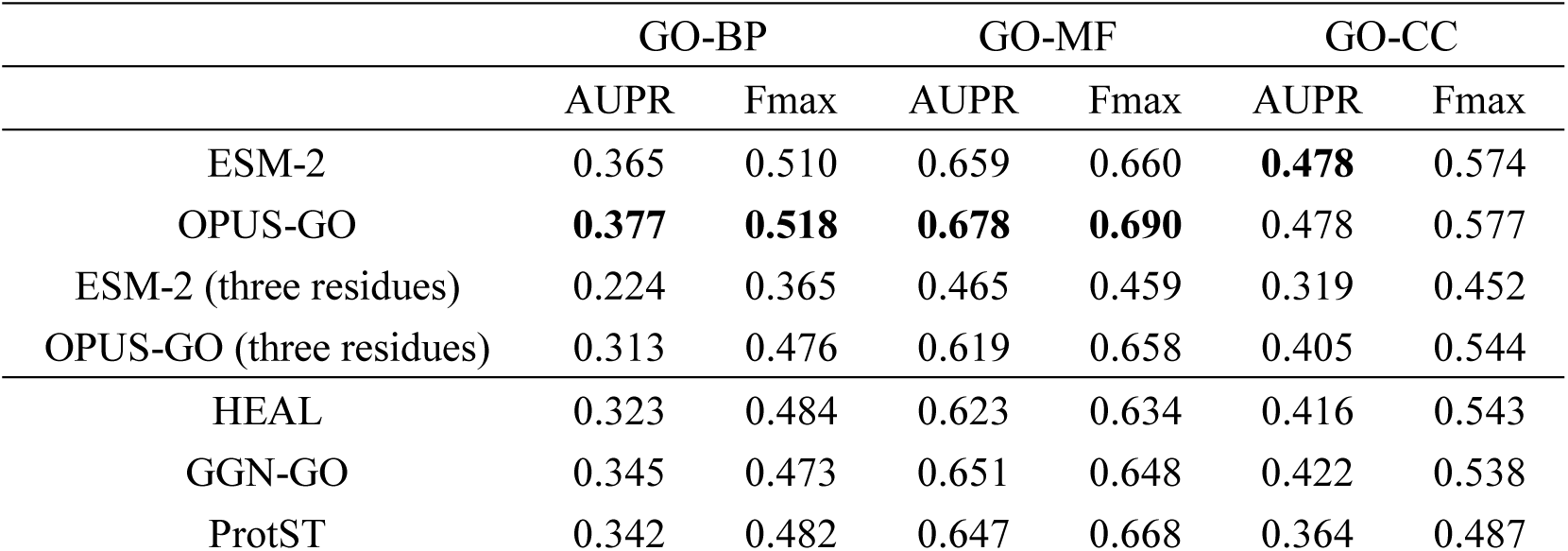
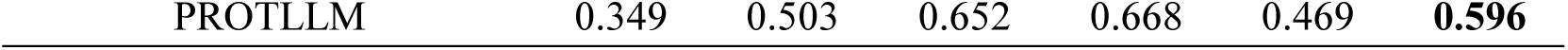
The results of different methods on three protein GO term prediction datasets. Best results for each metric are shown in boldface.

|  | GO-BP |  | GO-MF |  | GO-CC |  |
| --- | --- | --- | --- | --- | --- | --- |
|  | AUPR | Fmax | AUPR | Fmax | AUPR | Fmax |
| ESM-2 | 0.365 | 0.510 | 0.659 | 0.660 | **0.478** | 0.574 |
| OPUS-GO | **0.377** | **0.518** | **0.678** | **0.690** | 0.478 | 0.577 |
| ESM-2 (three residues) | 0.224 | 0.365 | 0.465 | 0.459 | 0.319 | 0.452 |
| OPUS-GO (three residues) | 0.313 | 0.476 | 0.619 | 0.658 | 0.405 | 0.544 |
| HEAL | 0.323 | 0.484 | 0.623 | 0.634 | 0.416 | 0.543 |
| GGN-GO | 0.345 | 0.473 | 0.651 | 0.648 | 0.422 | 0.538 |
| ProtST | 0.342 | 0.482 | 0.647 | 0.668 | 0.364 | 0.487 |

Furthermore, to validate the significance of the residues identified by OPUS-GO, we select three residues with the highest probability among the predictions for all labels, and use the average value of their ESM-2 features as the sequence-level representative to train a sequence-level classification model (OPUS-GO (three residues) in Table 1). For comparison, we employ the average ESM-2 features of three randomly selected residues as the sequence-level representative, and train the classification model under the same experimental settings (ESM-2 (three residues) in Table 1). The results indicate that the accuracies remain relatively stable when utilizing three residues selected by OPUS-GO, compared to those based on randomly selected residues. This demonstrates the effectiveness of OPUS-GO in locating residues that are associated with the corresponding label.

Due to the difficulty in associating many functional annotations with specific residues, we provide examples of annotations that can be directly linked to particular residues in Figure 2. These examples are based on the sequences whose 3D structures have been released in the Protein Data Bank (PDB). It is important to emphasize that OPUS-GO only utilizes the sequence of the target and does not require any structural information or information about other molecules, such as DNA, chemical molecules, or other protein partners. The results in Figure 2 demonstrate that OPUS-GO is capable of locating the residues associated with the corresponding labels with satisfactory accuracy.

**Figure 2.**
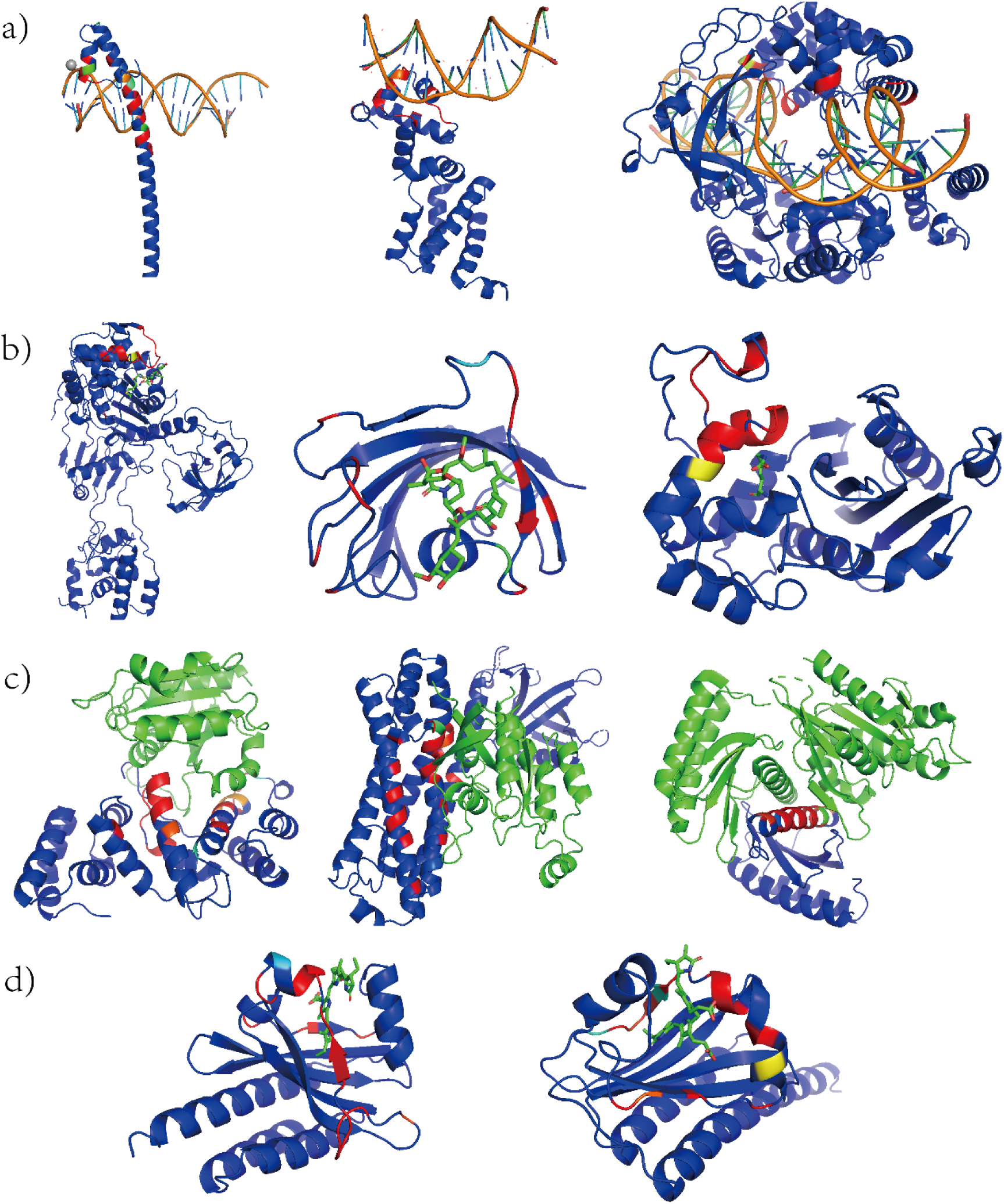
The interpretability results of OPUS-GO for protein GO term prediction. The results are visually represented using a color spectrum (ranging from blue to red) in PyMOL software, according to the predicted probability of corresponding label. It is important to note that, in each case, only the sequence information of the blue protein structure is utilized. The structures of these proteins, as well as the DNA in subfigure a), the chemical molecules in subfigures b) and d), and the green protein structure related to signal transduction, are not used in OPUS-GO and are only shown here for illustrative purposes. a) The probabilities of each residue for the label “DNA binding”. b) The probabilities of each residue for the label “drug binding”. c) The probabilities of each residue for the label “regulation of signal transduction”. d) The probabilities of each residue for the label “protein-chromophore linkage”.

For instance, Figure 2a showcases the results of OPUS-GO on three targets whose combined structure with DNA has been determined. The results indicate that OPUS-GO successfully identifies the residues related to the DNA binding site. Furthermore, Figure 2c presents the predicted probabilities of each residue for the label “regulation of signal transduction”. In the first case depicted on the left, the figure illustrates the structural basis for the activation of ARF GTPase (35). OPUS-GO employs the sequence of the Sec7 domain (depicted in blue) and the results demonstrate that it successfully identifies the residues (highlighted in red) on the Sec7 domain that are associated with its binding site with nucleotide-free ARF1 GTPase (depicted in green). Meanwhile, we present two examples of predicted probabilities for the label “proteolysis” in Figure S1. The results indicate that OPUS-GO successfully identifies the residues associated with the corresponding active sites (36, 37).

We also examine the results on the target for its various functional labels. Specifically, Figure 3a and Figure 3b present the predicted probabilities of each residue for the labels “transition metal ion binding” and “DNA binding”, respectively, on the same target. The findings indicate that OPUS-GO effectively distinguishes between residues pertinent to each functional group. Additionally, Figure 3c-f show the predicted probabilities of each residue corresponding to the labels “heme binding”, “tetrapyrrole binding”, “quinone binding” and “organic acid binding” on the same target. Despite belonging to distinct functional categories, it is noteworthy that heme is a subclass of tetrapyrrole, and quinone falls within the category of organic acids. The results reveal a significant similarity in the residues with high probabilities for “heme binding” and “tetrapyrrole binding”, as well as for “quinone binding” and “organic acid binding”, thereby confirming OPUS-GO’s ability to accurately pinpoint residues associated with their respective functions with good interpretability.

**Figure 3.**
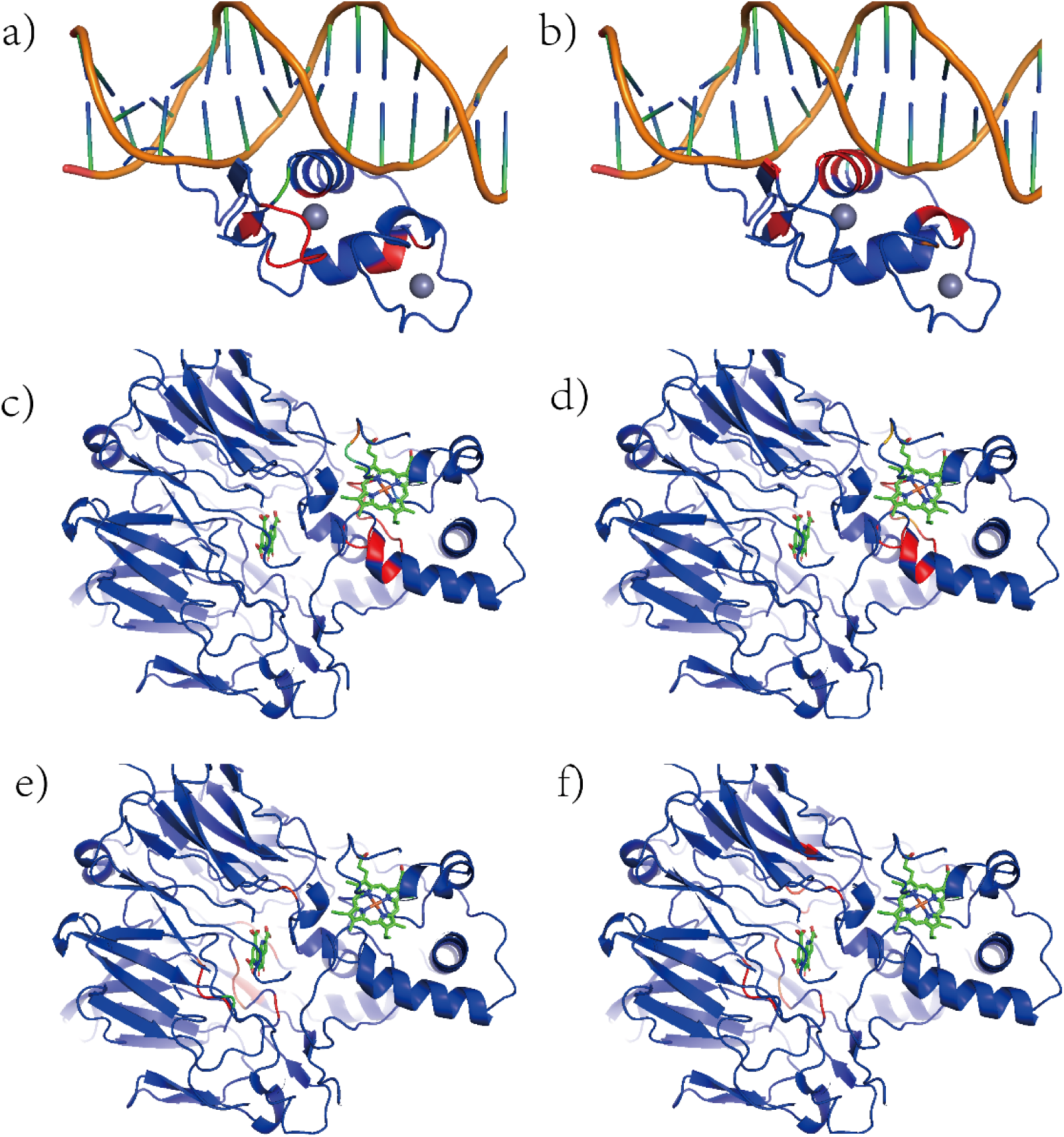
The interpretability results of OPUS-GO for protein GO term prediction. The results are visually represented using a color spectrum (ranging from blue to red) in PyMOL software, according to the predicted probability of corresponding label. a) The probabilities of each residue for the label “transition metal ion binding”. b) The probabilities of each residue for the label “DNA binding”. c) The probabilities of each residue for the label “heme binding”. d) The probabilities of each residue for the label “tetrapyrrole binding”. e) The probabilities of each residue for the label “quinone binding”. f) The probabilities of each residue for the label “organic acid binding”.

We compare the residue-level interpretability results of OPUS-GO and the ESM-2 using Grad-CAM, as well as GGN-GO (32), which also provides interpretability results based on Grad-CAM. In this context, we construct three test sets for the evaluation of the DNA binding, calcium ion binding, and heme binding, utilizing the targets in the GO-MF test set whose residue-level labels can be found from the BioLip database (38). These test sets consist of 34, 53, and 49 targets, respectively. Subsequently, we calculate the Area Under the ROC Curve (AUC) and F1-score for each method. Additionally, we introduce a shifted true positive criterion: if the binding site residue is located within 1 or 2 residues from the predicted residue, it is considered a true positive, and we compute the F1-shift-1 and F1-shift-2 metrics accordingly. As shown in Table 2, OPUS-GO consistently exhibits superior performance compared to the other two methods. We illustrate some examples of each method in Figure 4.

**Figure 4.**
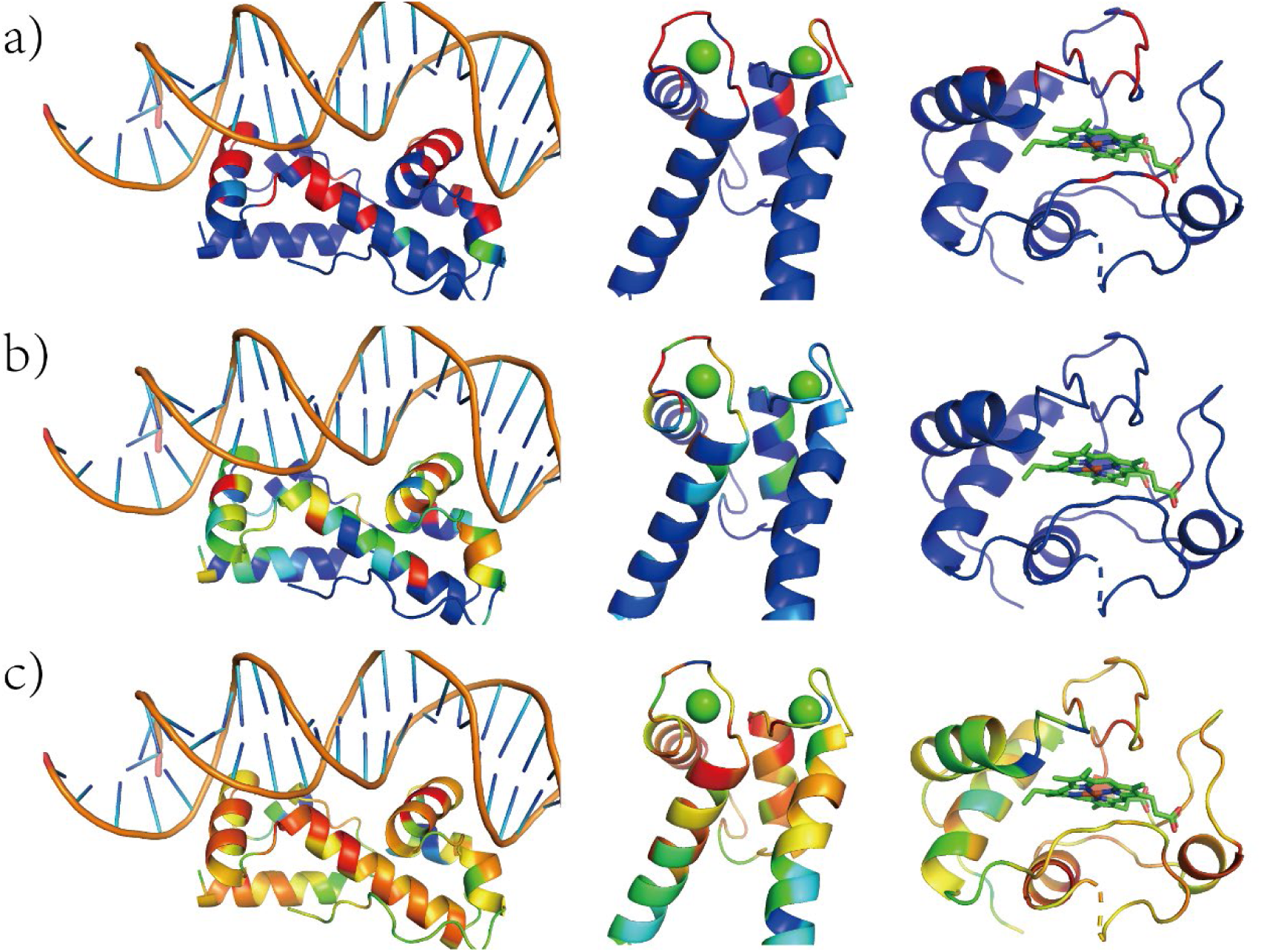
The interpretability results of each method. The results are visually represented using a color spectrum (ranging from blue to red) in PyMOL software, according to the predicted probability of corresponding label. a) The results of OPUS-GO. b) The results of ESM-2 with Grad-CAM. c) The results of GGN-GO.

**Table 2.**
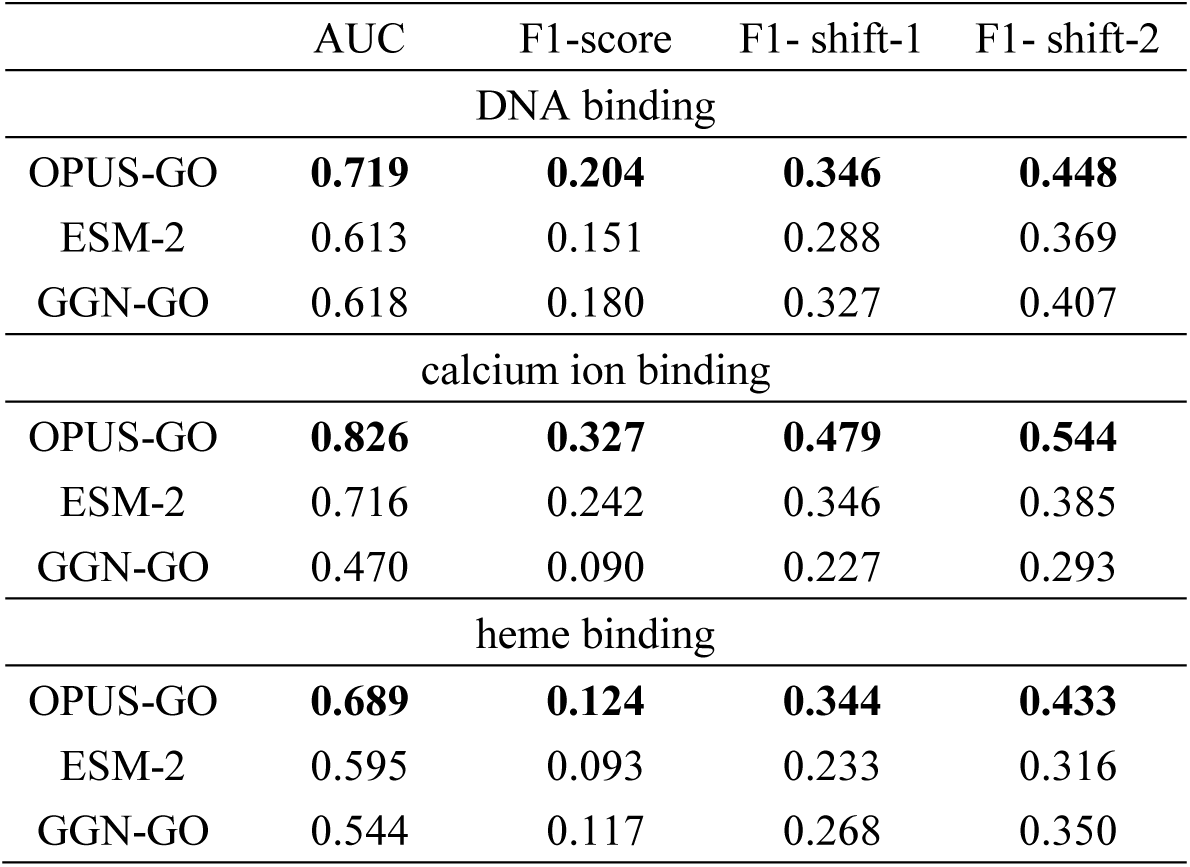
The residue-level interpretability results of different methods on three binding related datasets. Best results for each metric are shown in boldface.

|  | AUC | F1-score | F1- shift-1 | F1- shift-2 |
| --- | --- | --- | --- | --- |
| DNA binding |  |  |  |  |
| OPUS-GO | **0.719** | **0.204** | **0.346** | **0.448** |
| ESM-2 | 0.613 | 0.151 | 0.288 | 0.369 |
| GGN-GO | 0.618 | 0.180 | 0.327 | 0.407 |
| calcium ion binding |  |  |  |  |
| OPUS-GO | **0.826** | **0.327** | **0.479** | **0.544** |
| ESM-2 | 0.716 | 0.242 | 0.346 | 0.385 |
| GGN-GO | 0.470 | 0.090 | 0.227 | 0.293 |
| heme binding |  |  |  |  |
| OPUS-GO | **0.689** | **0.124** | **0.344** | **0.433** |
| ESM-2 | 0.595 | 0.093 | 0.233 | 0.316 |
| GGN-GO | 0.544 | 0.117 | 0.268 | 0.350 |

### Performance of OPUS-GO on protein location and ion binding prediction datasets

Protein localization prediction is a task aimed at determining the subcellular locations of proteins. In this study, we utilize two standard benchmarks from DeepLoc (39). The first benchmark involves subcellular localization prediction across 10 distinct locations, while the second benchmark involves binary localization prediction. We adhere to the official dataset splits for both tasks, with 10% of the training data being randomly selected to form the validation set. Furthermore, we incorporate the Metal Ion Binding task (40), which aims to predict the presence of metal ion-binding sites. All of them belong to the multi-class classification tasks, and the average accuracy in the test set are used as the evaluation metrics for each method.

As shown in Table 3, OPUS-GO exhibits superior sequence-level classification accuracy in all cases compared to the baseline method, which utilizes the average value of ESM-2 features across all residues as the sequence-level representative (ESM-2 in Table 3). Furthermore, the performance of a sequence-level classification model trained using the average value of three residues selected by OPUS-GO (OPUS-GO (three residues) in Table 3) surpasses that of the model trained using the average of all residues (ESM-2 in Table 3), highlighting OPUS-GO’s effectiveness in identifying residues associated with the corresponding label. In contrast, the model that relies on three randomly selected residues exhibits a significant decline in performance (ESM-2 (three residues) in Table 3). We present two examples of predicted probabilities for the Metal Ion Binding task in Figure S2. The results indicate that OPUS-GO successfully identifies the residues associated with the regions related to the metal ion binding sites.

**Table 3.** The results of different methods on protein location and ion binding prediction datasets. Best results for each metric are shown in boldface.

|  | Binary<br>Localization | Subcellular<br>Localization | Ion Binding |
| --- | --- | --- | --- |
| ESM-2 | 0.916 | 0.802 | 0.776 |
| OPUS-GO | 0.929 | **0.837** | 0.818 |
| ESM-2 (three residues) | 0.845 | 0.717 | 0.657 |
| OPUS-GO (three residues) | **0.931** | 0.834 | **0.827** |

It is noteworthy that, in contrast to the GO term prediction task presented in Table 1, where three residues may not suffice to cover all multiple labels, the tasks presented in Table 3 are multi-class classification tasks, with only one label for each sequence. Therefore, three residues may be sufficient to cover the particular label, leading to improved results.

### Performance of OPUS-GO on two protein EC number prediction datasets

The Enzyme Commission (EC) number is a widely employed system for categorizing protein enzyme functions through a standardized four-digit structure. This system provides a unified framework and accelerates the development in enzyme engineering. To assess the efficacy of our method on this task, we utilize two standard benchmarks: one proposed by DeepFRI, consisting of 538 labels (27, 30), and another recently introduced by GraphEC, comprising 5,106 labels (41). The former is denoted as EC in Table 4. The latter includes two test tests: the New-392 (42) and the Price-149 (43). We follow the official dataset splits for both benchmarks, allocating 10% of the training data, selected at random, to serve as the validation set for the second benchmark. Both benchmarks belong to the multi-label classification tasks, and we adopt Area Under the Precision-Recall Curve (AUPR) and Fmax as evaluation metrics for each method. Furthermore, for the second benchmark, we report the accuracy of the top-1 prediction for each method.

**Table 4.** The results of different methods on two protein EC number prediction datasets. Best results for each metric are shown in boldface.

|  | EC |  | NEW-392 |  |  | Price-149 |  |  |
| --- | --- | --- | --- | --- | --- | --- | --- | --- |
|  | AUPR | Fmax | AUPR | Fmax | TOP1 | AUPR | Fmax | TOP1 |
| ESM-2 | 0.899 | 0.862 | 0.485 | 0.610 | 0.543 | 0.219 | 0.439 | 0.396 |
| OPUS-GO | **0.902** | **0.881** | **0.487** | **0.641** | **0.566** | **0.244** | **0.448** | **0.423** |
| ESM-2 (three residues) | 0.610 | 0.597 | 0.222 | 0.369 | 0.347 | 0.038 | 0.144 | 0.114 |
| OPUS-GO (three residues) | 0.872 | 0.863 | 0.454 | 0.570 | 0.561 | 0.184 | 0.393 | 0.396 |
| GraphEC | - | - | 0.276 | 0.458 | 0.402 | 0.121 | 0.258 | 0.195 |
| CLEAN | - | - | - | - | 0.541 | - | - | 0.403 |

As shown in Table 4, OPUS-GO demonstrates better sequence-level classification accuracy in all cases when compared to the baseline methods. For the second benchmark, we incorporate the results of GraphEC (41) and CLEAN (42) for comparison, given that they utilized the same dataset in their studies. In GraphEC, for each residue, features derived from protein language model ProtTrans and structural predictions made by ESMFold are integrated through a geometric graph learning network to determine the final EC number. CLEAN (Contrastive Learning-Enabled Enzyme Annotation) is a machine learning algorithm designed to assign EC numbers to enzymes with enhanced accuracy, reliability, and sensitivity, leveraging features also obtained from protein language model ESM-1b. Since the probability of each label cannot be obtained from the output of CLEAN’s algorithm, we only calculate the accuracy of its prediction using the output label with the highest probability. In addition, for the NEW-392 dataset, we utilize only 391 results from GraphEC for calculation as it is unable to produce results for the target “Q694C5”. The results indicate that OPUS-GO outperforms both GraphEC and CLEAN. Consistent with the results observed in previous tasks, the performance of the sequence-level classification model trained using the average value of three residues selected by OPUS-GO (OPUS-GO (three residues) in Table 4) exceeds that of the model trained using the average of three randomly selected residues (ESM-2 (three residues) in Table 4), indicating the effectiveness of OPUS-GO in locating residues associated with the corresponding label.

The accuracy of the top-1 prediction for each method is presented in Figure 5a (NEW-392) and Figure 5b (Price-149) at four different levels. OPUS-GO consistently demonstrates superior performance compared to other methods. Additionally, Figure 5c-f displays the interpretability results of OPUS-GO for the protein EC number prediction task. Notably, although OPUS-GO solely utilizes the sequence information of the target protein, we have selected examples whose 3D structures have been released in the PDB for clearer presentation. The residues colored in red represent the active site of each target protein, while the yellow residues indicate positions that OPUS-GO predicts to be related to its enzymatic functional annotation. The results indicate that, in some examples (Figure 5c-e), the predicted positions are situated near the active sites. Conversely, there are also examples where OPUS-GO is unable to precisely identify the active sites (Figure 5f).

**Figure 5.**
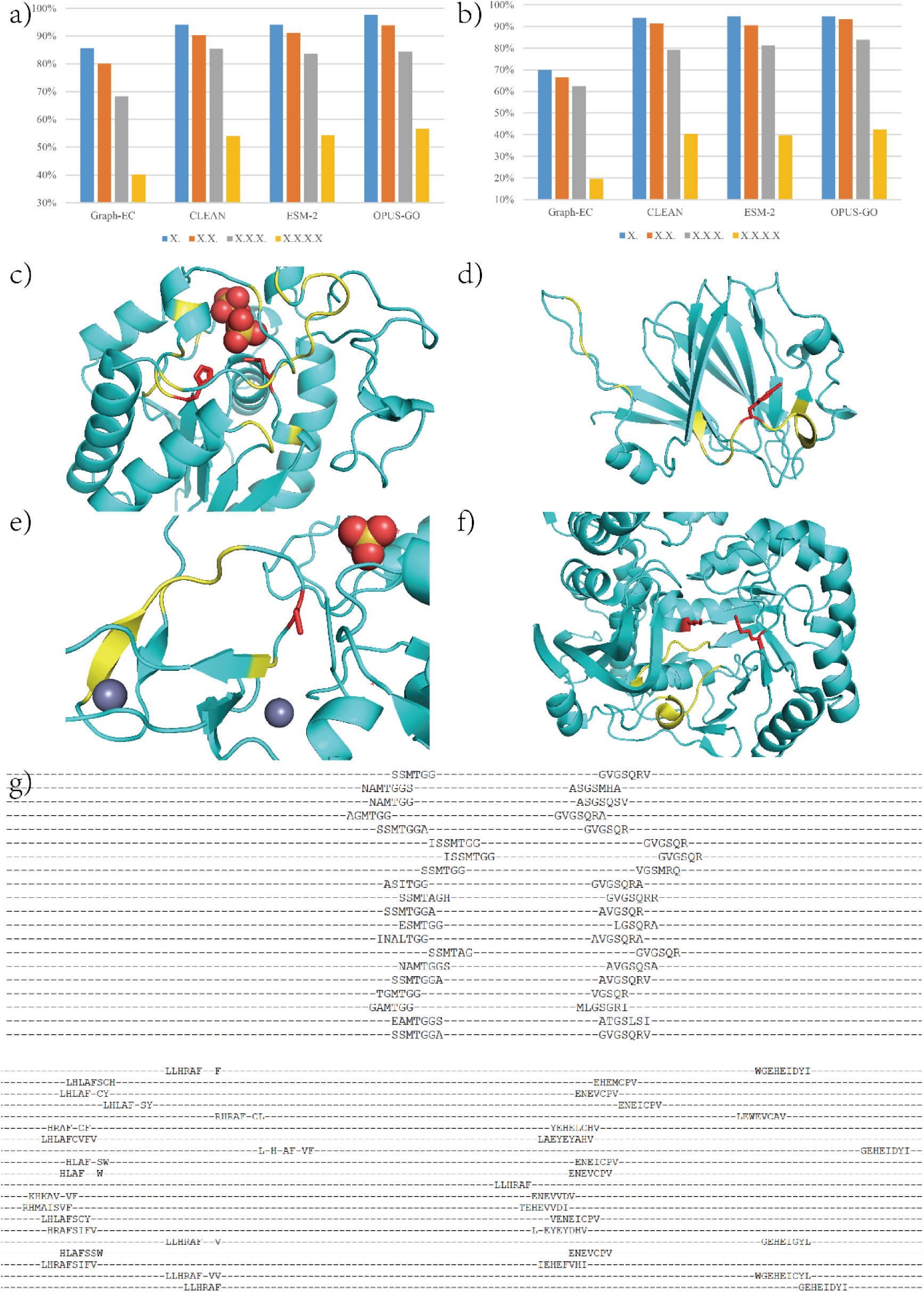
The results of OPUS-GO on protein EC number prediction task. a) The accuracy of the top-1 prediction for each method on NEW-392. b) The accuracy of the top-1 prediction for each method on Price-149. c-f) The interpretability results of OPUS-GO. Only the sequence information of the cyan protein structure is utilized in OPUS-GO. The residues with side chains illustrated and colored in red represent the active site of each target. The residues colored in yellow represent the positions that OPUS-GO predicts to be related to its enzymatic functional annotation. g) The residues that OPUS-GO identified as being related (with a probability greater than 0.5) to the label “5.3.3.2” (Isopentenyl-diphosphate Delta-isomerase) for the sequences within the training set.

However, it is important to note that the interpretability feature of OPUS-GO is not specifically tailored for identifying active sites. Instead, it identifies residues that it deems to be most directly associated with the corresponding labels, which may or may not align with the active sites. To verify this point, we conduct an analysis of the targets with the EC number “5.3.3.2” (Isopentenyl-diphosphate Delta-isomerase) within the training set, examining the residues that OPUS-GO identified as being most related to the label “5.3.3.2” for each sequence. As illustrated in Figure 5g, instead of identifying specific activate sites, OPUS-GO successfully identifies consistent patterns among these targets. These patterns can be categorized into two classes and are relatively conserved, and situated near the binding sites of each target. Considering the potential essentiality of these patterns for the function of the enzyme, OPUS-GO emerges as a valuable tool in protein design, offering crucial patterns that can facilitate the design of proteins with specific functionalities.

### Performance of OPUS-GO on RNA splice site prediction datasets

In eukaryotic genome annotation, splice sites (SS) denote the precise locations where introns are recognized and excised from pre-mRNA (precursor messenger RNA) by the spliceosome. These splice sites are indispensable for the accurate processing of genes and the subsequent synthesis of functional proteins. Specifically, there are two types of splice sites: the donor site, positioned at the exon-intron boundary, and the acceptor site, situated at the intron-exon junction (44). To assess the performance of our method, we employ the benchmark datasets proposed by Spliceator, which consists of sequences of 400 nucleotides in length (44). A subset of 10% of the training data is randomly selected to constitute the validation set. Since it is a binary classification task (splice site or non-splice site), we report the average accuracy and the Area Under the ROC Curve (AUC) for each method on the test set. For RNA downstream tasks, we utilize features extracted from two language models: RiNALMo (14) and ProtRNA (6).

As illustrated in Table S1, in RNA splice site prediction, utilizing the average value of features derived from RiNALMo (RiNALMo in Table S1) across all residues as the sequence-level representative, outperforms the use of features from ProtRNA (ProtRNA in Table S1). Meanwhile, OPUS-GO exhibits comparable sequence-level classification accuracy when employing features from RiNALMo (OPUS-GO (RiNALMo) in Table S1). Furthermore, when OPUS-GO relies on features from ProtRNA, it achieves significantly enhanced accuracy (OPUS-GO (ProtRNA) in Table S1).

More importantly, the dataset from Spliceator (44) positions the SS at the center of each sequence, enabling us to conduct a statistical evaluation. Consequently, we calculate the average probability of each residue being labeled as a “splice site” across all sequences in the test set. As shown in Figure 6, OPUS-GO effectively highlights the residues in the central regions when relying on both RiNALMo (Figure 6a-b) and ProtRNA (Figure 6c-d). Furthermore, the concentration of the highlights is more pronounced when utilizing features from the language model that performs better in this task.

**Figure 6.**
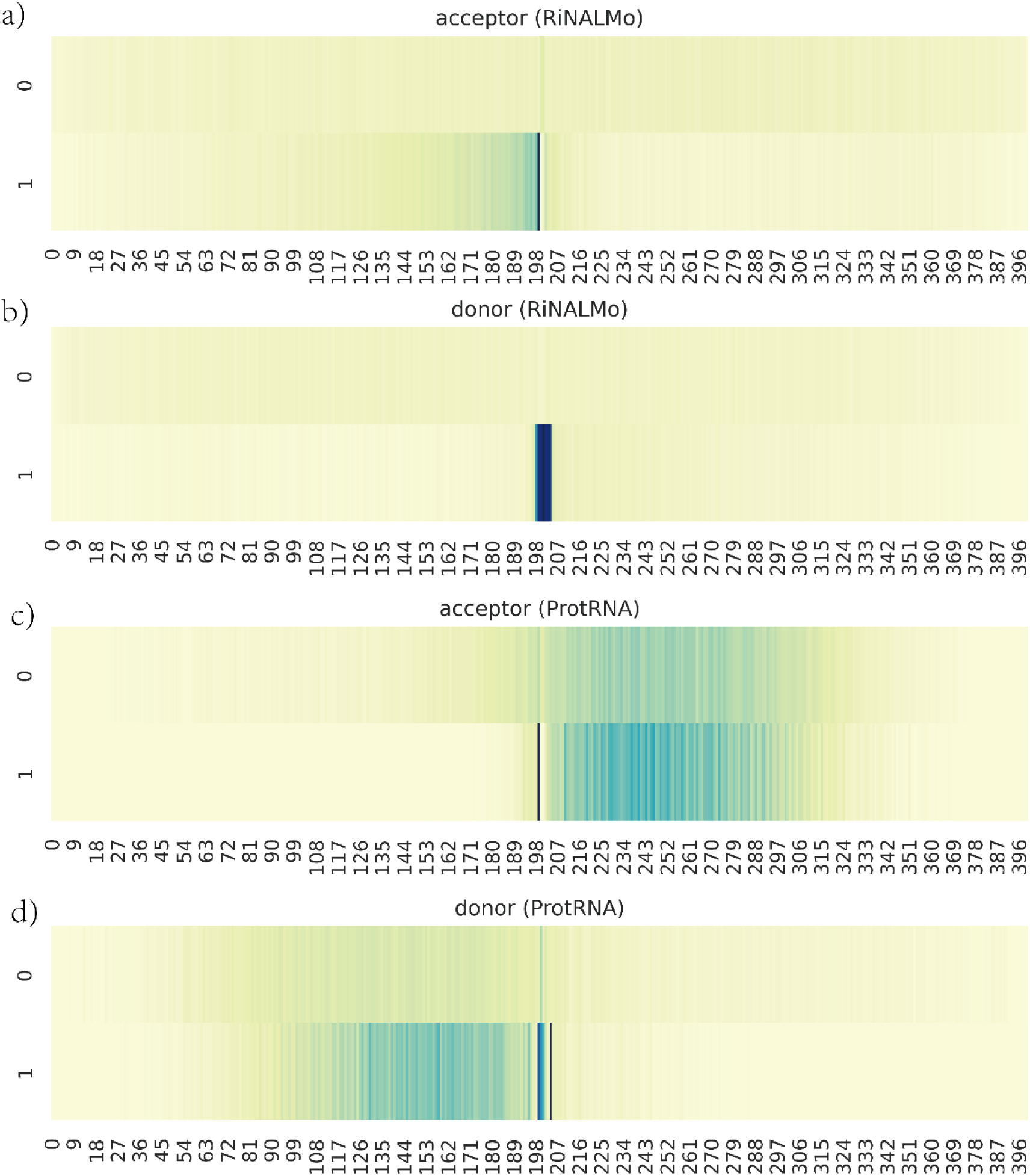
The results of OPUS-GO on RNA splice site prediction task. The average probability of each residue being labeled as a “splice site” across all sequences in the test set. “0” refers to the sequences without splice site, and “1” refers to the sequences containing splice site. a) The heatmap of acceptor site prediction using the features from RiNALMo. b) The heatmap of donor site prediction using the features from RiNALMo. c) The heatmap of acceptor site prediction using the features from ProtRNA. d) The heatmap of donor site prediction using the features from ProtRNA.

### Performance of OPUS-GO on Protein-RNA interaction prediction datasets

RNA binding proteins (RBPs) play a pivotal role in numerous biological processes, particularly in regulating RNA metabolism and function. Therefore, elucidating the binding profiles of RBPs is essential for comprehending their functional roles. In this task, we utilize three datasets regarding RBP binding sites collected by PrismNet (45). A validation set is constructed by randomly selecting 10% of the training data. Similar to the RNA splice site prediction task, the prediction of Protein-RNA interactions is a binary classification problem, involving the distinction between sequences that can bind to the target protein (“binding”) and those that cannot (“non-binding”). We present the average accuracy and the Area Under the ROC Curve (AUC) for each method evaluated on the test set.

As shown in Table 5, OPUS-GO demonstrates superior performance in terms of sequence-level classification accuracy in most cases when compared to the baseline methods. These baseline methods utilize the average value of features, derived from corresponding language model across all residues, as the representative for sequence-level classification.

**Table 5.**
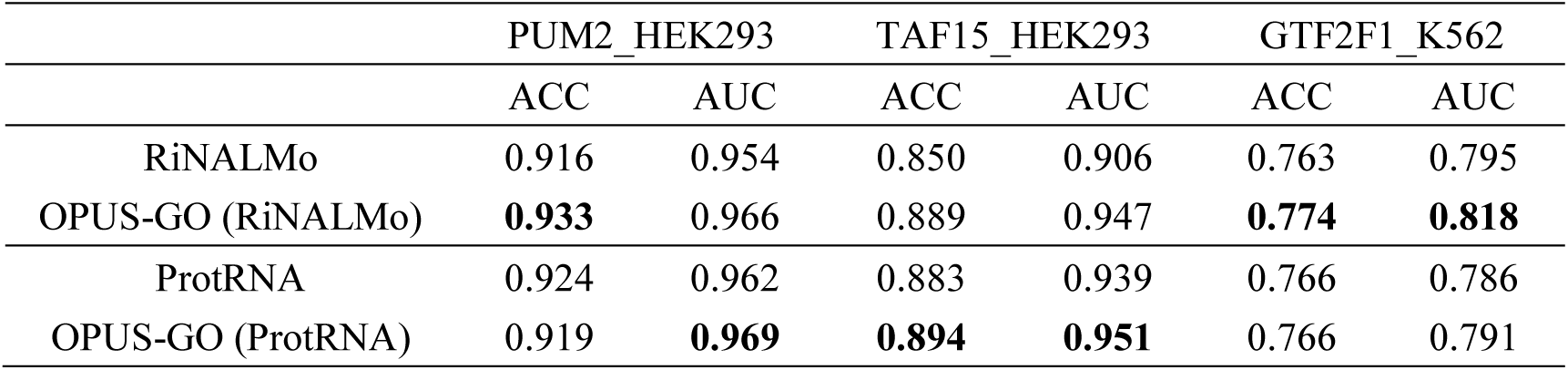
The results of different methods on three RBP binding sites prediction datasets. Best results for each metric are shown in boldface.

According to PrismNet, in these three datasets, the peak obtained from CLIP experiments is defined as the binding site for corresponding sequence. The length of each binding site is standardized to 101 nucleotides (nt). For regions shorter than 101 nt, the sequence is expanded symmetrically from the center to both sides, while for regions longer than 101 nt, the sequence is trimmed symmetrically from both sides to the center. This standardization allows us to conduct a statistical evaluation similar to that in RNA splice site prediction task. As shown in Figure S3, OPUS-GO successfully highlights the residues in the central regions utilizing both language models on the PUM2_HEK293 and TAF15_HEK293 datasets (Figure S3a-d). However, OPUS-GO exhibits a lack of effectiveness on the GTF2F1_K562 test set (Figure S3e-f). As illustrated in Table 5, the sequence-level classification results for the GTF2F1_K562 dataset are inferior to those obtained for the other two datasets. Since OPUS-GO relies on features derived from language models, the performance of these models directly influences the accuracy of OPUS-GO.

## Method

### Framework of OPUS-GO

OPUS-GO is built upon the Multiple Instance Learning (MIL) strategy and is well-suited for addressing multi-class or multi-label classification tasks. Specifically, it conceptualizes a sequence as a bag, with each residue’s feature within the sequence serving as an individual instance. Based on the sequence-level label of each sequence, we can train a residue-level classifier that is capable of classifying each residue according to the sequence-level label. Consequently, this classifier can be utilized to identify the most representative residues for each corresponding label.

The workflow of OPUS-GO is depicted in Figure 7. Initially, we extract features for each residue across each sequence target using relevant biological language models, such as ESM-2 (5), RiNALMo (14) and ProtRNA (6). Subsequently, a residue-level classifier is employed to predict the likelihood of each residue being associated with the corresponding sequence-level labels. The architecture of this classifier comprises a FeedForward module, a RMSNorm layer (24), and a Dropout Layer with a dropout rate of 0.5. The Swish activation function is employed within this architecture. No further aggregation of information from other residues in the sequence is necessary, as it has already been incorporated through the transformer block during the pretraining phase of the biological language models.

**Figure 7.**
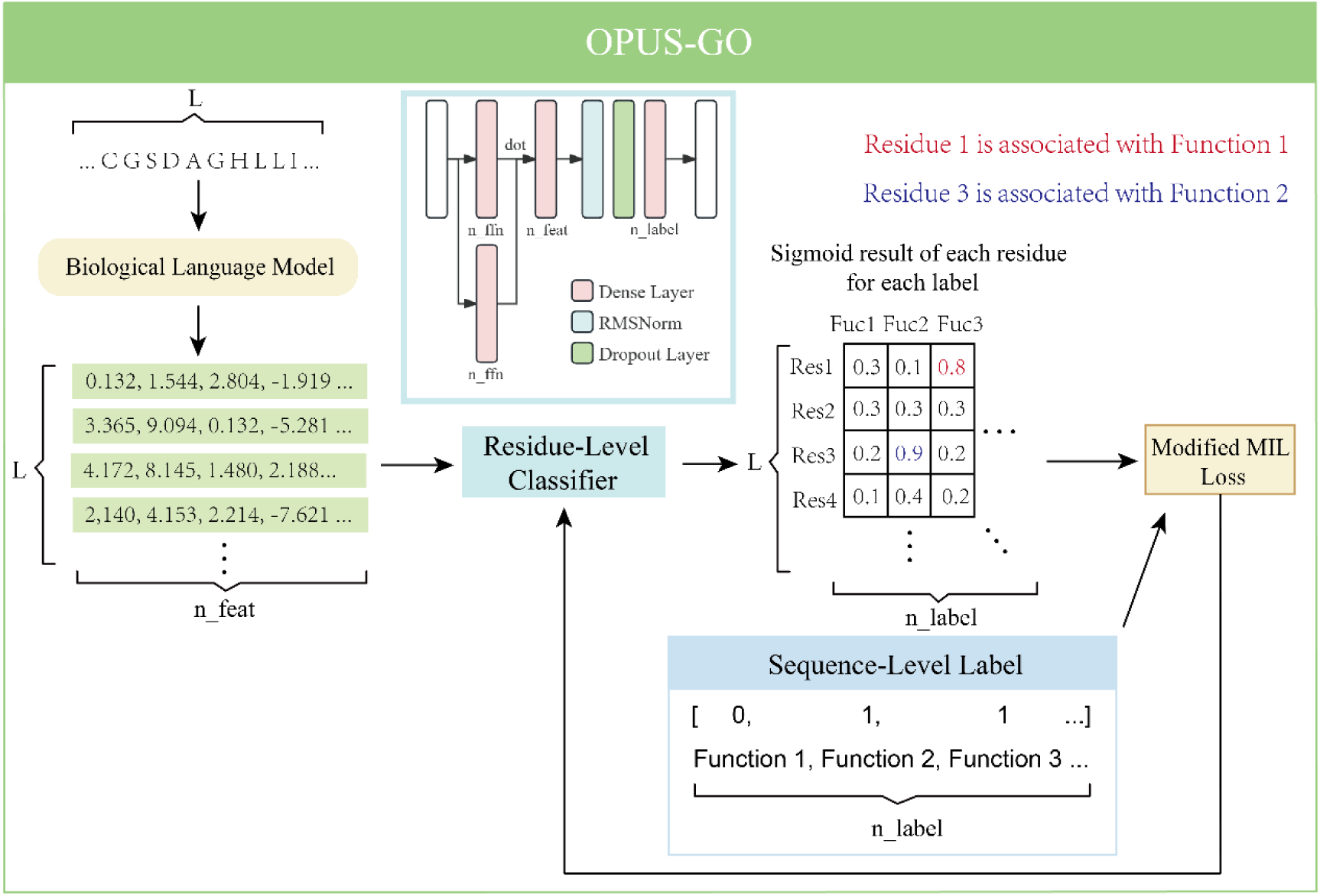
The workflow of OPUS-GO. “L” denotes the length of the protein/RNA sequence. “n_feat” denotes the number of features obtained from the biological language models. Specifically, the features utilized are derived from the last transformer block, such as the representations from the 33rd layer of ESM-2. “n_ffn” denotes the dimension of the hidden layers within the FeedForward module. “n_label” denotes the number of sequence-level labels, each treated as a binary classification task. The final output comprises the sigmoid values of each residue for each label. The sequence-level output for each label is determined by selecting the maximum sigmoid value across all residues within the corresponding column. Meanwhile, residues with a sigmoid value greater than 0.5 are considered to be associated with the respective label.

In this study, a modified MIL loss function is introduced to compute the loss and update the classifier. Since the sequence-level annotation may consists of multiple labels, the binary cross-entropy loss is applied to each label. The modified MIL loss can be further divided into two components. As shown in Algorithm S1, “n_top” is a hyperparameter that specifies the number of residues utilized in the calculation for each label. In OPUS-GO, we set “n_top” to the lesser value between 10% of all residues in sequence and 10 residues. For each positive label, we select the top “n_top” residues based on their probabilities in descending order and assign these residues with positive labels for loss computation. Concurrently, we randomly sample an equivalent number of residues from those not previously chosen and assign their labels as negative for loss computation.

Additionally, we introduce an auxiliary MIL loss, as detailed in Algorithm S2. This loss term exclusively utilizes negative samples in its calculation. The hyperparameter “n_top_labels” specifies the number of labels considered for computation. In OPUS-GO, if the total number of labels exceeds 5,000, “n_top_labels” is set to 20. Otherwise, all labels are considered. Firstly, the top “n_top_labels” labels are selected based on their maximum probabilities across all residues, sorted in descending order. Subsequently, for each negative label within the “n_top_labels” labels, the top “n_top” residues are chosen based on their probabilities, sorted in descending order, and assigned negative labels for loss computation. Both the auxiliary MIL loss and the previously mentioned MIL loss are added with equal weight to update the parameters of the residue-level classifier.

The incorporation of the hyperparameter “n_top_labels” is essential in tasks with an extremely large number of labels, such as those exceeding 5,000. In auxiliary MIL loss, it is observed that negative labels typically exhibit low probabilities for each residue. However, the accurate penalization of negative samples that exhibit relatively high probabilities for negative labels is crucial. Utilizing the average value across all negative labels would undermine the significance of pertinent information. Therefore, we select the “n_top_labels” based on their maximum probabilities across all residues, thereby ensuring that the necessary penalizations are adequately applied.

In this study, we set the hyperparameter “n_top_labels” with values of 1, 10, 20, 50, 200, 500, and 5,106 on the EC number prediction benchmark proposed by GraphEC (41). Subsequently, we train the models respectively. As shown in Table S2, the results suggest that the F1-Score attains its maximum value when “n_top_labels” is set to 20. Therefore, we recommend a default setting of 20 for tasks with an extremely large number of labels.

During the training phase, the Adam optimizer (46) is utilized. The model is trained for a maximum of 25 epochs, with an initial learning rate set to 1e-3. This learning rate is halved when a decrease in validation accuracy is observed after each epoch. Early stopping is implemented if the learning rate has been reduced four times. OPUS-GO is developed using TensorFlow (47) version 2.4 and trained on four NVIDIA Tesla V100 GPUs. The total batch size is set to 16.

During the inference phase, the sequence-level prediction is aggregated by selecting the maximum value for each label across all residues, with residues possessing a sigmoid value greater than 0.5 being considered as associated with the corresponding label. For a given label, its probability is assigned as the maximum sigmoid value among all residues. Additionally, for multi-class classification tasks, where each sequence has only one label, the final output is determined by selecting the label with the highest sigmoid value.

### Data and Software Availability

The training and evaluation codes and pre-trained models of OPUS-GO for each downstream task can be downloaded from http://github.com/thuxugang/opus_go.

## Conclusion

As the volume of biological data expands rapidly, the development of large-scale biological language models (BLMs) has been significantly accelerated, resulting in impressive performance across various scenarios and substantially enhancing the accuracy of numerous downstream tasks. In computational biology, existing BLMs-based methods often prioritize enhancing sequence-level classification accuracy while neglecting the significance of residue-level interpretability. This primarily arises from the relative ease of acquiring sequence-level annotations compared to the challenges associated with obtaining residue-level annotations. However, interpretability remains a crucial issue, as it holds the potential to offer valuable biological insights that transcend mere accuracy improvements. In this study, we introduce OPUS-GO, a method that exclusively utilizes sequence-level annotations to provide both sequence-level and residue-level classification results, thereby enabling the identification of residues most critical for the sequence-level annotation. Our results demonstrate that, based on the features derived from BLMs and out modified Multiple Instance Learning (MIL) strategy, OPUS-GO shows superior performance in sequence-level classification compared to baseline methods for most of protein and RNA downstream tasks. Additionally, OPUS-GO offers strong interpretability by accurately pinpointing residues associated with respective labels. Furthermore, OPUS-GO’s effectiveness on datasets containing over 5,000 labels suggests its widespread applicability in various contexts.

In contrast to MIL, which not only offers robust interpretability but also has the potential to concurrently enhance model performance, Class Activation Map (CAM)-based methods are restricted to use as a post-training analysis tool, providing the interpretability of model without affecting its accuracy. On the other hand, based on the MIL framework, an instance-level classifier can be obtained and utilized independently of its surrounding context, unlike GAM-based methods that require context-specific application. In summary, both strategies possess their unique advantages. While many methods utilize Grad-CAM to showcase the interpretability, this study proposes an alternative approach using MIL to address this issue. Our results indicate that OPUS-GO offers superior residue-level interpretability compared to CAM-based methods. Several potential enhancements to MIL-based still exist, such as in OPUS-GO, where we simply utilize the maximum sigmoid value among all residues as the final result for a given label. However, this approach may be susceptible to outliers. A more delicate aggregation strategy may further improve the sequence-level accuracy.

It should be noted that OPUS-GO is capable of identifying residues that it deems to be most directly related to the corresponding labels, which may or may not correspond to a specific site of interest, such as active sites in the EC number prediction task. However, the results indicate that, rather than identifying specific active sites, OPUS-GO successfully detects consistent patterns within each category of enzymes. Given the potential significance of these patterns for enzyme function, OPUS-GO emerges as a valuable tool in protein design, offering crucial patterns that can facilitate the design of proteins with specific functionalities.

## Acknowledgements

J.M. wants to thank the support from the Science and Technology Innovation Plan of Shanghai Science and Technology Commission (No. 23JS1400200), and the Research Fund for International Senior Scientists (No. W2431060). G.X. wants to thank the support from the National Natural Science Foundation of China (No. 32300535).

## Declaration of interests

Q.W. is an employee of Harcam Biomedicines who is bound by confidentiality agreements that prevent her from disclosing the competing interests in this work. The other authors declare no competing interests.

## Author Contributions

G.X., Q.W., and J.M. conceived the work. G.X. designed the algorithm and implemented the software. G.X., Y.L., R.Z., and X.X. collected the data and evaluated the results on the various downstream tasks. All authors participated in discussion of the project and writing of the manuscript.

